# Development of an environmental DNA method for monitoring fish communities: ground truthing in diverse lakes with characterised fish faunas

**DOI:** 10.1101/394718

**Authors:** Jianlong Li, Tristan W. Hatton-Ellis, Lori-Jayne Lawson Handley, Helen S. Kimbell, Marco Benucci, Graeme Peirson, Bernd Hänfling

## Abstract

1. Accurate, cost-effective monitoring of fish is required to assess the quality of lakes under the European Water Framework Directive (WFD). Recent studies have shown that environmental DNA (eDNA) metabarcoding is an effective and non-invasive method, which can provide semi-quantitative information on fish communities in large lakes.
2. This study further investigated the potential of eDNA metabarcoding as a tool for WFD status assessment by collecting and analysing water samples from eight Welsh lakes and six meres in Cheshire, England, with well described fish faunas. Water samples (*N* = 252) were assayed using two mitochondrial DNA regions (Cytb and 12S rRNA).
3. eDNA sampling indicated the presence of very similar species in the lakes compared to those expected on the basis of existing and historical information. In total, 24 species were detected with a total of 111 species occurrences in the lakes studied using eDNA. Secondly, there was a significant positive correlation between expected faunas and eDNA data in terms of confidence of species occurrence (Spearman’s *r* = 0.74, *df* = 109, *p* <; 0.001). Thirdly, eDNA data can estimate relative abundance with the standard five-level classification scale (“DAFOR”). Lastly, four ecological fish communities were characterised using eDNA data which agrees with the pre-defined lake types according to environmental characteristics.
4. *Synthesis and applications*. This study provides further evidence that eDNA metabarcoding could be a powerful and non-invasive monitoring tool for WFD purpose in a wide range of lake types, considerably outperforming other methods for community level analysis.

## Introduction

The impact of anthropogenic pressures on aquatic ecosystems is ubiquitous and usually leads to alterations of the environment (Scheffer et al. 2001). To mitigate negative human effects, the first step in the improvement of the ecological quality of degraded water bodies is the development of an assessment system to evaluate the current situation. The main European initiative for water quality assessment and improvements, the Water Framework Directive 2000/60/EC (WFD), requires all member states to reach “good” ecological status of lakes, rivers and ground waters based on biological elements including phytoplankton, macrophytes and phytobenthos, benthic invertebrates and fish (CEC 2000).

Fish are widely considered as relevant for detecting and quantifying impacts of anthropogenic pressures on lakes and reservoirs (Argillier et al. 2013). Although most UK WFD ecological tools have been developed and deployed, an effective fish-based assessment tool for lakes remains problematic (Winfield 2002; Kelly et al. 2012). The current fish-based indices (e.g. Argillier et al. 2013) or tools (e.g. “FIL2” in Kelly et al. 2012) rely on semi-quantitative capture-based methods such as electro-fishing or gill-netting to provide information on fish composition and abundance, as well as age-structure of fish populations. Furthermore, the choice of survey methods is heavily dependent on the depth and size of lake. However, both of these sampling methods are relatively laborious and thus expensive, sometimes destructive, taxonomically biased, cannot be deployed in all situations (e.g. in or near dense vegetation, or entire water column in a very deep lake) and have poor sampling accuracy and precision. These drawbacks restrict their ability to meet WFD requirements (Kubecka et al. 2009; Winfield et al. 2009). A future strategy for WFD purposes thus requires a highly cost-effective approach, with a minimum amount of destructive sampling.

A significant “game-changer” in biodiversity monitoring in recent years is the analysis of environmental DNA (eDNA) which is a non-invasive genetic method that takes advantage of intracellular or extra-organismal DNA in the environment to detect the presence of organisms (Lawson Handley 2015; Thomsen & Willerslev 2015; Taberlet et al. 2018). A particularly promising approach is to simultaneously screen whole communities of organisms using eDNA metabarcoding. Several studies have shown that eDNA metabarcoding using High-Throughput Sequencing offers tremendous potential as a complementary method tool to established monitoring methods for ecology and conservation of aquatic species (e.g. Hänfling et al. 2016; Port et al. 2016; Valentini et al. 2016). After reviewing and discussing the potential of eDNA metabarcoding as a tool for WFD status assessment, Hering et al. (2018) suggested that this approach is well-suited for fish biodiversity assessment considering representativeness, sensitivity, precision, comparability, cost-effectiveness, and environmental impact. The mitochondrial COI gene is widely, but not unanimously accepted as the standard metabarcode for metazoans (e.g. Riaz et al. 2011; Clarke et al. 2014). However, with environmental samples, most primers targeting the COI region amplify large proportions of microbial species (e.g. Stat et al. 2017). This fact remains the strongest reason for the use of other genes such as mitochondrial cytochrome b (Cytb) and 12S rRNA (12S) regions. A prototype eDNA tool assayed by Cytb and 12S regions for fish biodiversity assessment was tested in three lakes in the English Lake District, and initial results from this are promising (Hänfling et al. 2016). However, in order to understand factors such as responses to ecological pressures, taxon specific biases, sampling requirements in different lake types and management of low confident occurrence, a much larger dataset from a range of lakes with different ecological characteristics is required. Thus, the objectives of this study were (a) to broaden the lake fish dataset using eDNA-based metabarcoding analysis by collecting and analysing water samples from 14 UK lakes with well described fish faunas; (b) to explore a possible approach to evaluate the confidence of species presence based on site occupancy and read counts and (c) to use a relative abundance scale to estimate species abundance by comparing the eDNA metabarcoding results to historical data.

## Materials and methods

### Study sites

The distribution of the 14 lakes including eight Welsh lakes and six Cheshire meres are shown in Figure 1, and the general characteristics are outlined in Appendix S1 Table S1.1 based on data from the UK Lakes Portal (Hughes et al. 2004). In brief, the 14 lakes can be divided into three types according to environmental characteristics including Type 1: three low alkalinity lakes that are broadly upland in character (“CWE” Llyn Cwellyn, “PAD” Llyn Padarn and “OGW” Llyn Ogwen); Type 2: two high alkalinity but shallow lakes (“PEN” Llyn Penrhyn and “TRA” Llyn Traffwll) on the west coast of Anglesey (North Wales) which are close to sea and accessible for migratory fish, and Type 3: nine high alkalinity but shallow lakes that are dominated by coarse fish (“KEN” Kenfig Pool, “LLB” Llan Bwch-llyn, “LLG” Llangorse Lake, “MAP” Maer Pool, “CAM” Chapel Mere, “OSS” Oss Mere, “FEN” Fenemere, “WLF” Watch Lane Flash and “BET” Betley Mere) (Appendix S2).

**Figure 1.**
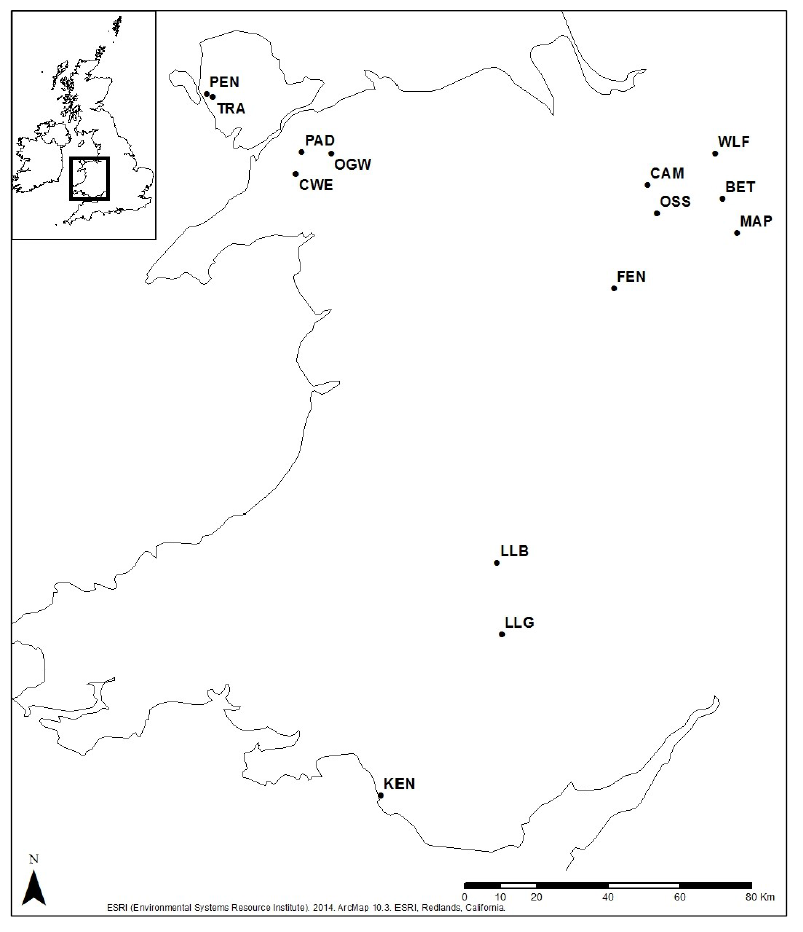
Distribution of sampling lakes. Sampling lake codes: (CWE) Llyn Cwellyn; (PAD) Llyn Padarn; (OGW) Llyn Ogwen; (PEN) Llyn Penrhyn; (TRA) Llyn Traffwll; (KEN) Kenfig Pool; (LLB) Llan Bwch-llyn; (LLG) Llangorse Lake; (MAP) Maer Pool; (CAM) Chapel Mere; (OSS) Oss Mere; (FEN) Fenemere; (WLF) Watch Lane Flash; (BET) Betley Mere. The GPS coordinates of sampling locations in each lake are listed in Appendices S3 & S4.

### eDNA capture, extraction, library preparation and sequencing

#### eDNA capture and extraction

In total, 252 samples were collected from the 14 lakes between 01/12/2015 to 12/07/2016 (Appendix S1 Table S1.2). The detail of sampling strategy can be found in Appendix S5.1. All samples were filtered within 24 hrs of collection. Two litres of each water sample were filtered through a 47 mm diameter 0.45-µm mixed cellulose acetate and nitrate filter (Whatman, UK) using Nalgene filtration units in combination with a vacuum pump (15–20 in. Hg; Pall Corporation, USA). Our previous study demonstrated that the 0.45-µm filters are suitable for fish metabarcoding, with low variation and high repeatability between the filtration replicates (Li et al. 2018a) compared to other filtration methods. The filtration units were cleaned with 10% v/v commercial bleach solution (containing ~3% sodium hypochlorite) and 5% v/v microsol detergent (Anachem, UK), and then rinsed thoroughly with de-ionised water after each filtration run to prevent cross-contamination. For each sampling lake, a filtration blank (2 L deionised water) was filtered before sample filtration in order to test for possible contamination at the filtration stage. After filtration, all filters were placed into 50 mm sterile petri dishes using sterile tweezers, sealed with parafilm and then immediately stored at –20°C until extraction. DNA extraction was carried out using the DNeasy PowerWater Kit (QIAGEN, Germany) following the manufacturer’s protocol. The DNA was eluted in 100 μL 10 mM Tris (Solution PW6) and stored at –20 °C freezer.

#### Library preparation and sequencing

Sequencing libraries were generated from PCR amplicons targeting Cytb and 12S. We previously tested both fragments *in vitro* on 22 UK freshwater fish species and *in situ* on three deep lakes in the English Lake District, and demonstrated their suitability for eDNA metabarcoding of UK lake fish communities (Hänfling et al. 2016).

To enable the detection of possible PCR contamination, we included no-template controls (NTCs, i.e. negative control) of molecular grade water (Fisher Scientific, UK) and single-template controls (STCs, i.e. positive control) of cichlid fish DNA (the Eastern happy, *Astatotilapia calliptera,* a cichlid from Lake Malawi, which is not present in natural waters in the UK) within each library. All lake samples (*N* = 252), together with 14 sampling blanks, 14 filtration blanks, 14 NTCs, and 14 STCs were included in each library construction (*N* = 308) for sequencing on an Illumina MiSeq. Three PCR technical replicates were performed for each sample then pooled to minimise bias in individual PCRs. All PCRs were set up using eight-strip PCR tubes in a PCR workstation with UV hood and HEPA filter in the eDNA laboratory at University of Hull (UoH) to minimise the risk of contamination.

The Cytb locus, targeting a 414-bp vertebrate-specific fragment, was amplified with the L14912 and H15149 primer pair (Kocher et al. 1989) using a one-step library preparation protocol (Kozich et al. 2013) (see more detail in Appendix S5.2).. To improve clustering during initial sequencing, the final 10 pM denatured Cytb library mixed with 30% PhiX control was sequenced with the MiSeq reagent kit v3 (2×300 cycles) at UoH. The 12S locus targets a 106-bp vertebrate-specific fragment amplified with the 12S_V5_F and 12S_V5_R primer pair (Riaz et al. 2011). Following the consistently lower sequencing yield of the one-step protocol for the 12S compared to the Cytb in previous studies, we decided to switch to a two-step library preparation protocol using nested tagging (Kitson et al. 2018) (see more detail in Appendix S5.2). Since the heterogeneity spacers included in the two-step library preparation protocol will improve the nucleotide diversity, the denatured 12S library (13 pM) was mixed with lower amount of PhiX control (i.e. 10%) compared to the Cytb library, then sequenced with the MiSeq reagent kit v2 (2×250 cycles) at UoH.

### Data analysis

#### Collation of fish data

Existing fish data for the 14 lakes was collated using a range of data sources (see the detail in Appendix S2). These included site-specific surveys carried out by Natural Resources Wales or the Environment Agency and their predecessor bodies, third party fishery surveys, and data published in the literature and *ad hoc* records stored on databases such as the National Biodiversity Network Atlas (https://nbnatlas.org). Based on existence of survey data and expert opinion (TWH-E and GP), fish species were placed into four general categories with confidence of species presence/absence for each lake (Table 1), reflecting both the available data and the uncertainty around it. Assessing relative abundance *a priori* was more challenging. For the Welsh lakes, an abundance category approach was used, with each species being placed on DAFOR scale (D = Dominant; A = Abundant; F = Frequent; O = Occasional; R = Rare) (Tansley 1993) using available data where possible and/or TWH-E’s expert opinion. All records where no abundance was recorded were assumed to be rare. For Cheshire meres, summaries of fish community composition and fish density per unit area (ind. ha^-1^ and kg ha^-1^) were produced from the Ecological Consultancy Ltd (ECON) fishery survey using point abundance sampling by electro-fishing (PASE) and seine netting in the same period as water samples collection with this study in 2016 (Appendix S1 Tables S1.3 & S1.4). The ECON fishery survey in 2016 was undertaken by ECON and commissioned by Natural England as part of an investigation of the impacts of fisheries on Sites of Special Scientific Interests (SSSIs) condition.

**Table 1.**
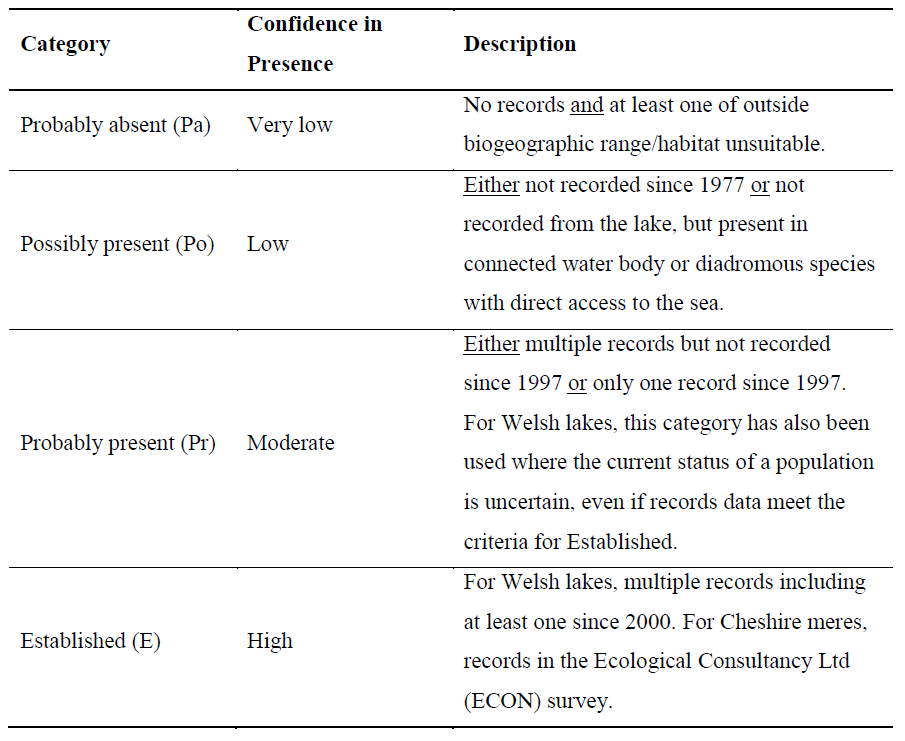
Criteria for assessing the confidence of species presence/absence using existing data.

| Category | Confidence in Presence | Description |
| --- | --- | --- |
| Probably absent (Pa) | Very low | No records <u>and</u> at least one of outside biogeographic range/habitat unsuitable. |
| Possibly present (Po) | Low | <u>Either</u> not recorded since 1977 <u>or</u> not recorded from the lake, but present in connected water body or diadromous species with direct access to the sea. |
| Probably present (Pr) | Moderate | <u>Either</u> multiple records but not recorded since 1997 <u>or</u> only one record since 1997. For Welsh lakes, this category has also been used where the current status of a population is uncertain, even if records data meet the criteria for Established. |
| Established (E) | High | For Welsh lakes, multiple records including at least one since 2000. For Cheshire meres, records in the Ecological Consultancy Ltd (ECON) survey. |

#### Bioinformatics analysis

Raw read data from Illumina MiSeq sequencing have been submitted to NCBI (BioProject: PRJNA454866). Bioinformatics analysis was implemented following a custom reproducible metabarcoding pipeline (metaBEAT v0.97.9) with custom-made UK freshwater fish reference databases (Cytb and 12S) as described in our previous study (Hänfling et al. 2016). Sequences which the best BLAST hit had a bit score below 80 or had less than 95%/100% identity (Cytb/12S) identity to any sequence in the curated databases were considered non-target sequences. The different level of similarity for each locus depended on the length of metabarcode and the knowledge of intraspecific diversity of the studied taxon. To assure full reproducibility of our bioinformatics analysis, the custom reference databases and the Jupyter notebooks for data processing have been deposited in an additional dedicated GitHub repository (https://github.com/HullUnibioinformatics/Li_et_al_2018_eDNA_fish_monitoring). The Jupyter notebook also performs demultiplexing of the indexed barcodes added in the first PCR reactions.

#### Low-frequency noise threshold

Filtered data were summarised into the number of sequence reads per species/sample for downstream analyses (Appendixes S3 & S4). After bioinformatics analysis, the low-frequency noise threshold (proportion of STC species read counts in the lake sample) was set to 0.07% and 0.3% for Cytb and 12S respectively to filter high-quality annotated reads passing the previous filtering steps that have high-confidence BLAST matches but may result from contamination during the library construction process or sequencing (De Barba et al. 2014; Hänfling et al. 2016; Port et al. 2016). The low-frequency noise threshold was guided by the analysis of sequence data from STCs under several thresholds. Filtered data were summarised in two ways for downstream analyses: (a) the number of sequence reads per species at each lake (hereafter referred to as read counts) and (b) the proportion of sampling locations in which a given species was detected (hereafter referred to as site occupancy).

#### Estimating abundance with eDNA

Based on our previous results (Hänfling et al. 2016), site occupancy is a better proxy for estimates of abundance than read counts. However, read counts should not be completely ignored because they still contain important abundance information. The maximum site occupancy across both loci was used as a score for species abundance (Table 2). In addition to site occupancy score, the maximum relative read count (i.e. proportion of read counts per species at each sampling lake) across both loci, and the number of loci with which the species was detected were used as confidence indicators to assign fish species into the same general categories which were used for non-eDNA survey data (Table 3). Those species that were assigned the lowest confidence scores (probably absent) were excluded from downstream analysis (i.e. roach *Rutilus rutilus* in CWE, bream *Abramis brama* and bullhead *Cottus gobio* in OGW, pike *Esox lucius* in PEN, bullhead in TRA, gudgeon *Gobio gobio* in BET). Each retained species was assigned a corrected abundance score by multiplying site occupancy score × confidence score. The corrected abundance score was used to assign each species to a relative abundance DAFOR scale (Table 4) (For instance, bream in LLB: 2 × 5 = 10 resulting in “Frequent” category in DAFOR scale).

**Table 2.** Site occupancy score based on maximum site occupancy across Cytb and 12S.

| Site occupancy score | Maximum site occupancy (across Cytb and 12S) |
| --- | --- |
| 1 | $> 0$ and $\leq 0.1$ |
| 2 | $> 0.1$ and $\leq 0.3$ |
| 3 | $> 0.3$ and $\leq 0.6$ |
| 4 | $> 0.6$ and $\leq 0.8$ |
| 5 | $> 0.8$ |

**Table 3.**
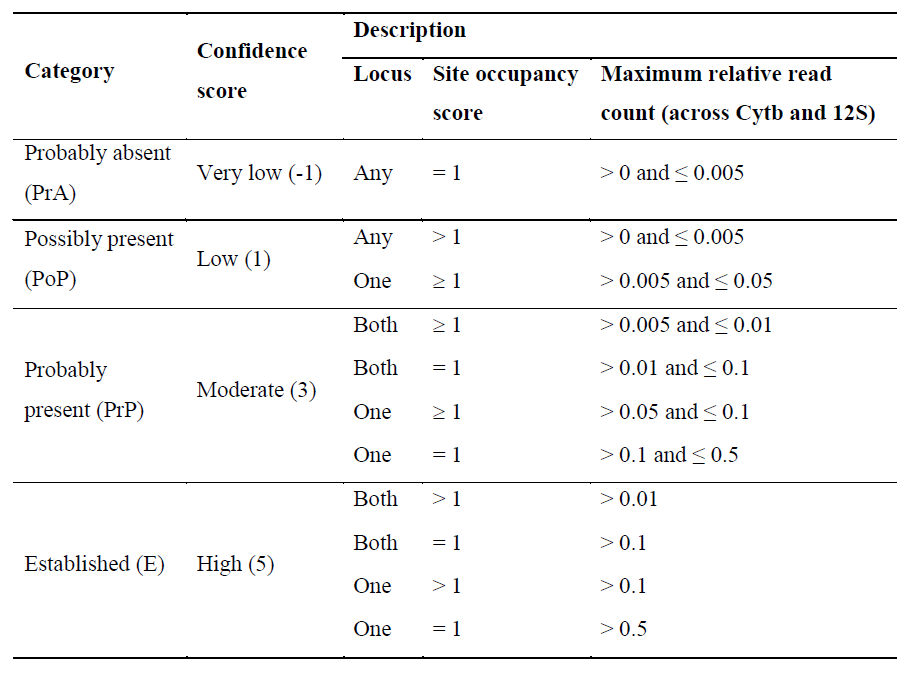
Criteria for assessing confidence of species occurrence using eDNA data.

**Table 4.** The relative abundance DAFOR scale based on eDNA data.

| Abundance rank<br>number | DAFOR scale | Corrected abundance<br>score (site occupancy<br>score × confidence<br>score) |
| --- | --- | --- |
| 0 | None | = 0 |
| 1 | Rare | $\geq 1$ and < 4 |
| 2 | Occasional | $\geq 4$ and < 9 |
| 3 | Frequent | $\geq 9$ and < 15 |
| 4 | Abundant | $\geq 15$ and < 25 |
| 5 | Dominant | = 25 |

#### Ecological and statistical analyses

All downstream analyses were performed in R v3.3.2 (R_Core_Team 2016) and graphs plotted using GGPLOT2 v2.2.1 (Wickham & Chang 2016). Before investigating species detection and abundance estimate with eDNA, we first evaluated whether Cytb and 12S datasets produced consistent results by calculating the Pearson’s product-moment correlation coefficient for both read counts and site occupancy. Spearman’s rank correlations were performed between historical data and eDNA data – firstly in terms of confidence of species presence and secondly in terms of relative abundance. To investigate differences in fish communities between sampling lakes, hierarchical clustering dendrogram was used to assess the existence of distinct community types using the function *fviz_dend* in FACTOEXTRA v1.0.4 (Kassambara & Mundt 2017) to extract and visualize the results calculated from the function *hclust* using Canberra distances method based on site occupancy. Furthermore, non-metric multidimensional scaling (NMDS) based on read counts in individual sampling location of lakes, allied with analysis of similarities (ANOSIM), were performed using the abundance-based Bray-Curtis dissimilarity index with the function *metaMDS* and *anosim* respectively in VEGAN v2.4-4 (Oksanen et al. 2017). The sequencing depth and sample-based rarefaction was performed using the *rarecurve* and *rarc* function in VEGAN v2.4-4 (Oksanen et al. 2017), respectively. The full R script is available on the GitHub repository (https://github.com/HullUnibioinformatics/Li_et_al_2018_eDNA_fish_monitoring/tree/master/R_script)

## Results

The Cytb and 12S libraries generated 17.15 and 15.97 million reads, respectively, with 15.91 and 13.4 million reads passing filters (including 49.01% or 11.07% PhiX). After quality filtering and removal of chimeric sequences, the average read count of lake sample was 12,503 for Cytb and 21,703 for 12S. After BLAST searches for taxonomic assignment, 59.47% ± 35.56% and 26.46% ± 19.60% reads in each lake sample were assigned to fish for Cytb and 12S, respectively. Rarefaction analysis revealed our sequencing depth to be sufficient to identify all of the amplified fish species present in the 14 lakes for both loci (Appendix S1 Figure S1.1).

Significant correlations were found between site occupancy and average read counts, for both loci and all sampling lakes apart from OGW (Pearson’s *r* = 0.81, *df* = 4, *p* = 0.05) and LLB (Pearson’s *r* = 0.94, *df* = 2, *p* = 0.06) for Cytb (Appendix S1 Figure S1.2a, b). Data from the two loci were significantly correlated (Pearson’s *r* consistently *p* <; 0.05) for site occupancy for each lake (Appendix S1 Figure S1.2c), and for average read counts apart from in three lakes: PAD (Pearson’s *r* = 0.55, *df* = 7, *p* = 0.12), TRA (Pearson’s *r* = 0.59, *df* = 5, *p* = 0.16) and LLB (Pearson’s *r* = 0.89, *df* = 2, *p* = 0.11) (Appendix S1 Figure S1.2d). The more detail of comparisons between these two loci in detection probabilities can be found in Appendix S5.3.

### Comparison of eDNA data and expected results

In total, 24 species with 111 species occurrences in 14 lakes were detected across Cytb and 12S. Perch *Perca fluviatilis* was the most common species which were detected in all lakes. The second most frequently occurring species was roach with detections in 10 lakes. In addition to these two species, pike were found in all nine coarse fish lakes (Table 5; Figure 2; Appendix S1 Figure S1.3). There was a significant positive correlation between expected faunas and eDNA data in terms of confidence of species occurrence (Spearman’s *r* = 0.74, *df* = 109, *p* <; 0.001). A total of 73 “established” species occurrences have been recorded across all lakes according to historical data. Of these, 72 (98.6%) were detected with eDNA (Table 5). The only false negative in the eDNA datasets was tench *Tinca tinca* in PEN. Tench were detected by 12S in PEN but at a read count less than the low-frequency noise threshold (Appendix S4; site occupancy = 0.3, average read counts = 3.9). Tench are likely to be very rare in PEN: only two tench were recorded by the 2016 Royal Society for the Protection of Birds survey and small numbers have also been detected in previous years (Appendix S2.4). Twenty-five of the 111 occurrences were “probably or possibly present”, and 13 were “very low confidence present” (i.e. probably absent records based on historical data) (Table 5).

There were consistent positive correlations between DAFOR scale based on eDNA data and expected DAFOR scale based on both historical data for Welsh lakes and fish density per unit area from the ECON survey of Cheshire meres (Figure 3). The correlations were significant in three out of eight Welsh lakes (PAD, OGW and KEN). Significant correlations were observed in three (MAP, CAM and BET) and four (MAP, CAM, FEN and WLF) Cheshire meres based on individual density (ind. ha^-1^) and biomass density (kg ha^-1^), respectively (Figure 3; Appendix S1 Figure S1.4).

Furthermore, the rigorous sampling of shore and offshore locations is carried out to compare the suitability of the different spatial sampling location (Appendix S1 Table S1.2). The more detail of comparisons between shore and offshore samples can be found in Appendix S5.4.

### Characterisation of fish assemblages using eDNA data

The hierarchical clustering dendrograms based on site occupancy for two loci indicated that there were four distinct community types compared to three pre-defined lake types according to environmental characteristics. Specifically, the clustering dendrograms accorded with the pre-defined lake Type 1 (CWE, OGW and PAD) and Type 2 (TRA and PEN), but indicated that WLF and BET can be divided from the pre-defined lake Type 3 (Figure 4a1, b1). The NMDS ordination allied with ANOSIM based on read counts for two loci confirmed the clustering dendrograms results that there were four distinct community types according to the predominant groups of fish including Community 1: salmonids and minnow *Phoxinus phoxinus*; Community 2: mixed diadromous fish; Community 3: coarse fish, and Community 4: bream-dominated coarse fish (Figure 4a2, b2). The *R* statistic in the ANOSIM global test with these four different communities were high (Appendix S1 Table S1.5, 0.75 ± 0.02) supporting statistical differences between communities (Appendix S1 Table S1.5, *p* = 0.001). The overall distance pattern of these four fish communities was that both coarse fish communities (Communities 3 & 4) were close to each other; the mixed diadromous fish community (Community 2) was between the coarse fish communities and the salmonids and minnow community (Community 1) (Figure 4; Appendix S1 Table S1.5).

The salmonids and minnow community (Community 1) is characteristic of three low alkalinity upland lakes (CWE, OGW and PAD). These sites were generally dominated by brown trout *Salmo trutta*, rainbow trout *Oncorhynchus mykiss* or Arctic charr *Salvelinus alpinus*, but also included minnow and Atlantic salmon *Salmo salar* (Figure 2; Appendix S1 Figure S1.3). CWE and PAD are designated as SSSIs under the Wildlife and Countryside Act 1981 (as amended) to conserve Arctic charr, and CWE, a oligotrophic lake, is also protected as a Special Areas of Conservation under the EU Habitats Directive. The eDNA data showed that PAD contained a small number of Arctic charr, whereas this species was dominant in CWE (Figure 2; Appendix S1 Figure S1.3). Rainbow trout were only detected in OGW, where they are regularly stocked by the local angling club (Appendix S2.3). Some of these sites may also contain low density of other species; usually diadromous species were detected using eDNA as well such as European eel *Anguilla anguilla* in all three upland lakes and three-spined stickleback *Gasterosteus aculeatus* in PAD. Perch have recently been recorded from PAD for the first time using captured-based survey methods (Appendix S2.2). Surprisingly, this species was detected using eDNA (though shown to be rare), in all of these three lakes (Figure 2; Table 5). The diadromous fish community (Community 2) is characteristic TRA and PEN which were dominated by European eel and three-spined stickleback. Other species included brown trout, nine-spined stickleback *Pungitius pungitius*, roach, perch and rudd *Scardinius erythrophthalmus* (Figure 2). The coarse fish communities consisted of bream, common carp *Cyprinus carpio*, pike, perch, roach, rudd and tench, plus some additional species (e.g. European eel, bullhead, three-spined stickleback and gudgeon). Pike, perch, roach and tench were dominant in KEN, MAP, CAM, LLB, LLG, OSS and FEN (Community 3); however, the dominant species in WLF and BET (Community 4) was bream (Figure 2; Appendix S1 Figure S1.3).

**Table 5.**
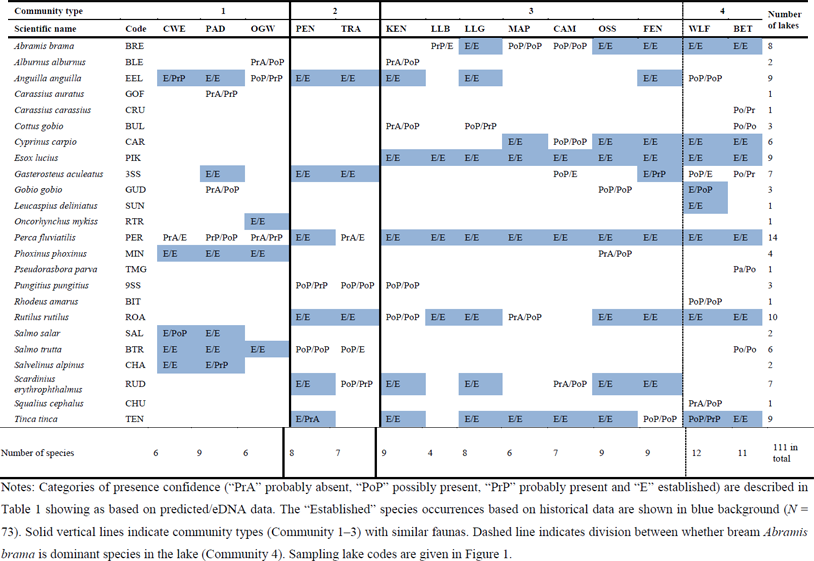
Correspondence of confidence of species occurrence between predicted and eDNA data in 14 lakes.

**Figure 2.**
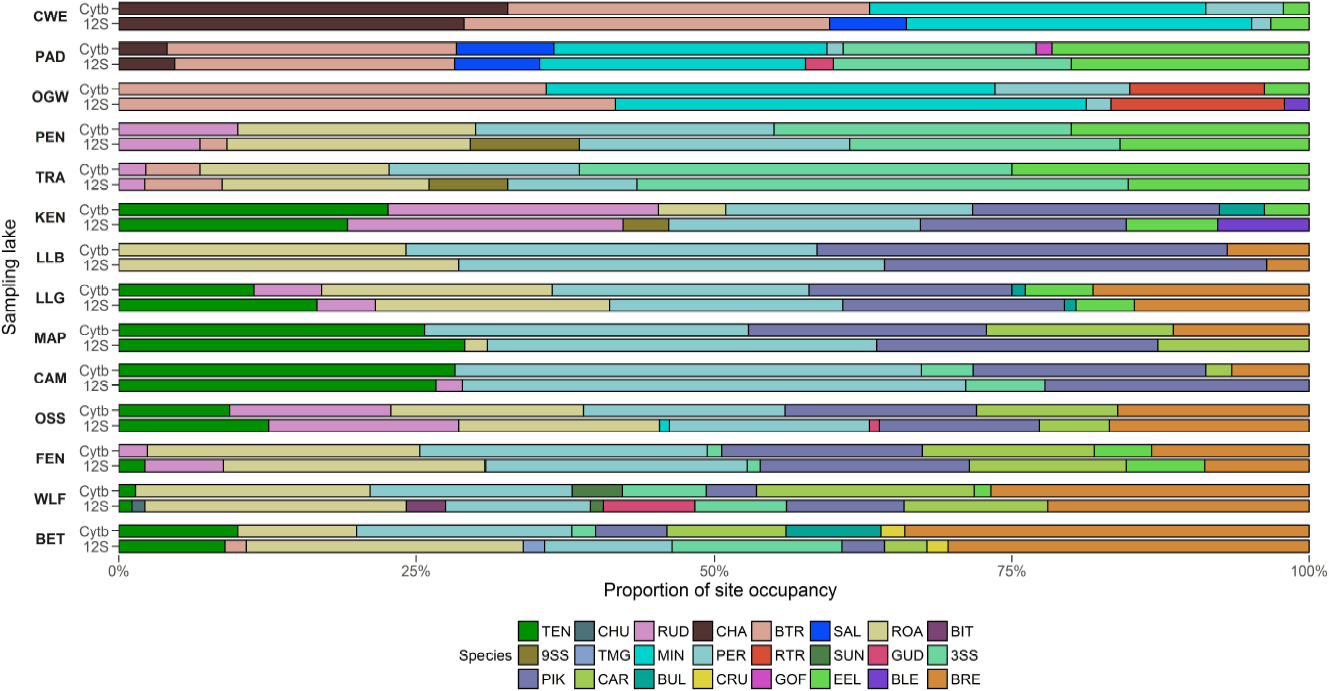
Species composition of site occupancy for Cytb and 12S. Species three letter codes and sampling lake codes are given in Table 5 and Figure 1, respectively.

**Figure 3.**
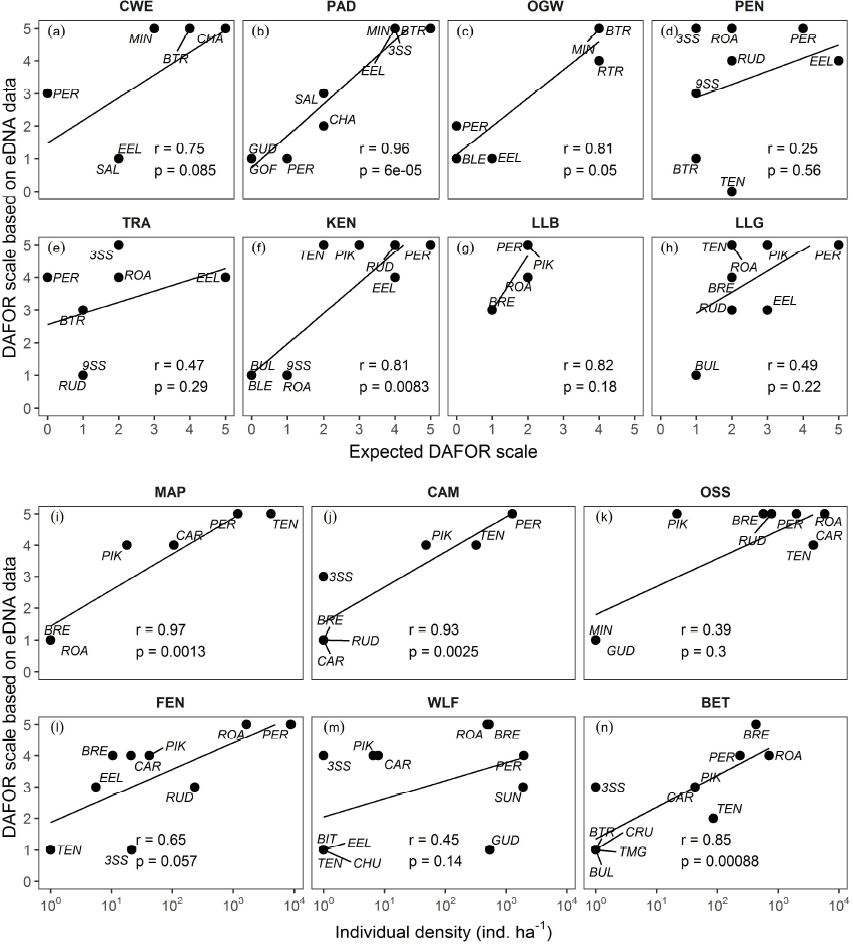
Correlations between DAFOR scale based on eDNA data and expected DAFOR scale based on existing data of Welsh lakes or individual density from the ECON survey of Cheshire meres. Sampling lake codes, abundance rank numbers to DAFOR scale and species three letter codes are given in Figure 1, Table 4 and Table 5, respectively.

**Figure 4.**
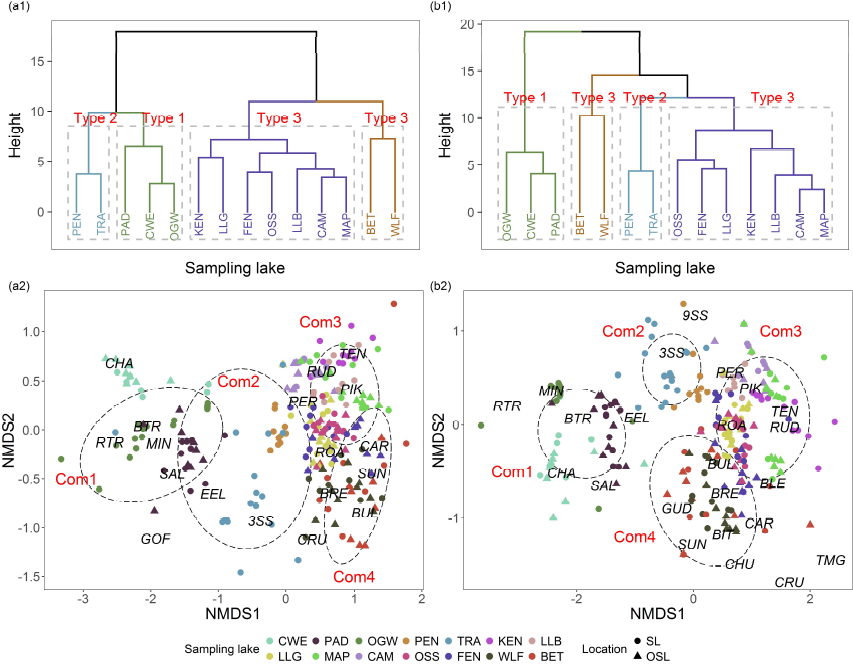
Hierarchical clustering dendrograms and non-metric multidimensional scaling (NMDS) ordination of the sampling lakes. Dendrograms using Canberra distances method based on site occupancy for (a1) Cytb and (b1) 12S. The dashed frames are drawn around each cluster in dendrograms. The three pre-defined lake types according to environmental characteristics. NMDS in individual sampling location of lakes using Bray-Curtis dissimilarity index based on read counts for (a2) Cytb and (b2) 12S. Each symbol corresponds to a sampling lake, with circles corresponding to shore samples (“SL”) and triangles corresponding to offshore samples (“OSL”) in NMDS ordination. The ellipse indicates 70% similarity level within each community type in ordinations. Scores for species taxa are plotted on the same axes to better visualise the ordination in space between species and samples. Species three letter codes and community type (Com 1–4) with individual lake codes are given in Table 5 and Figure 1, respectively.

## Discussion

This study confirmed key results from our previous study in that there was (a) a consistent, strong correlation between Cytb and 12S in terms of read counts and site occupancy; (b) a consistent, strong correlation between site occupancy and read counts and (c) site occupancy was consistently better than read counts for estimating relative abundance, for both Cytb and 12S datasets (Hänfling et al. 2016).

### Comparison of eDNA and existing data for species detection

A total of 73 “established” species occurrences were recorded across the 14 lakes according to existing and historical fish data. Of these, 72 (98.6%) were detected with eDNA, which demonstrated that eDNA metabarcoding outperformed the conventional surveys and observations using a range of sampling methods over date ranges spanning several decades in species richness estimates. This result is consistent with previous studies that have demonstrated that eDNA metabarcoding produced a more comprehensive species list than alternative survey techniques with a similar effort in both marine and freshwater ecosystems (Hänfling et al. 2016; Port et al. 2016; Valentini et al. 2016).

Moreover, there was a significant positive correlation between historical data and eDNA data in terms of confidence of species presence (Spearman’s *r* = 0.74, *df* = 109, *p* <; 0.001). Species occurrences that were assessed as “probably or possibly present” (25 of the 111 species occurrences) and “probably absent” (13 of the 111 species occurrences) based on distribution data or anecdotal evidence also fitted the eDNA criteria for lowered confidence levels. In most cases, the “probably absent” occurrences were at very low site occupancy (≤ 0.1) and read counts (~0.5%), just above our threshold for accepting a positive record. These records could be genuine detections of species that have previously been missed. For example, perch were recorded with both Cytb and 12S in CWE, OGW and TRA. The confidence of species present was either “established present” or “probably present” according to eDNA data. The recent appearance of this species in nearby lakes PAD and PEN (see detail in Appendix S2) suggest that the species is spreading in North Wales either by natural means or as a result of illegal introductions. In addition, the other “probably absent” occurrences could be either false positives from sequencing error, laboratory or environmental contamination. For instance, bleak *Alburnus alburnus* in OGW, goldfish *Carassius auratus* and Gudgeon in PAD, rudd in CAM, roach in MAP, topmouth gudgeon *Pseudorasbora parva* in BET and chub *Squalius cephalus* in WLF are most likely explained by low-level cross contamination or sequencing barcode misassignment since these species were only detected by either Cytb or 12S in single sample of the lake.

Barcode misassignment and tag jumps have been proved in metabarcoding studies (e.g. Deakin et al. 2014; Schnell et al. 2015). To minimise contamination or false assignments, further work needs to be done to optimise each step, as well as the incorporation of robust controls to identify contamination when it occurs. Together with a growing number of studies (Stat et al. 2017; Li et al. 2018b), the results demonstrate the importance of using two (or more) markers for metabarcoding to avoid problems due to primer bias (Clarke et al. 2014), gene copy number, PCR or sequencing artefacts (Schloss et al. 2011) and/or contamination. Other strategies such as using consistency of presence across technical replicates as used by Port et al. (2016) might be a more suitable approach to control for false positives if rare species are of particular interest. Where verification of rare species detections is considered a high priority, additional confirmation from targeted species approaches (e.g. quantitative PCR) and/or field surveys may be required.

### Use of eDNA for assessing relative abundance of fish

WFD specifies that abundance should be considered when determining ecological status; hence, current WFD approaches include estimates of abundance (often as abundance classes). For instance, a five-level scale, adapted from DAFOR scale, is accepted for Austrian standard and national monitoring techniques applied under WFD for surveying aquatic macrophytes (Pall & Moser 2009). Here, we used DAFOR scale to estimate fish relative abundance in the present study based on DNA-based identification to facilitate integration into current WFD approaches.

There were consistently positive correlations in terms of relative abundance between the eDNA data and historical data. However, these correlations were not always statistically significant. The correlations were not significant in TRA and PEN probably because threespined stickleback were more abundant based on eDNA data than expected. This species is often under-represented or overlooked in surveys using established fish capture methods (Hänfling et al. 2016). Moreover, tench were not detected by eDNA in PEN. LLB is a species-poor lake with only four species detected, which could reduce the statistical power. The non-significant correlation in WLF could be attributed to under-representation of threespined stickleback in previous fish surveys and reduced detection probability of gudgeon with Cytb. Although these results are generally encouraging, further work is critical to obtain enough statistical power to directly test the relationship between abundance estimates from eDNA and surveys using other methods. It is also important to investigate taxon-specific detection probabilities and abundance estimates so that a pressure-sensitive tool can be developed.

### Using eDNA to describe ecological communities

According to environmental characteristics, there were three pre-defined lake types. Encouragingly, four distinct community types could be identified based on clustering dendrograms and NMDS ordinations using eDNA data. Basically, the Community 1 and Community 2 agreed with the pre-defined lake Type 1 and Type 2, respectively. The predefined coarse fish lakes (Type 3) can be further divided into Community 3 and Community 4 based on whether bream was dominant species in the lake. These findings indicated that eDNA metabarcoding has great potential as fish-based assessment tool for WFD lake status assessment.

Specifically, our results showed that the low alkalinity lakes within a predominantly upland catchment were mainly dominated by salmonids reflecting their relatively deep and oligotrophic nature. Except from salmonids, all these sites contained minnow, most likely introduced by anglers using them as live bait (Hatton-Ellis 2005). Compared to the salmonids and minnow lakes (Community 1), perch, pike and cyprinids (such as bream, common carp, roach and tench) were prevalent in the coarse fish lakes (Communities 3 & 4) reflecting their relatively shallow and eutrophic nature. These findings support that eDNA profiles are suitable to reflect the eutrophic conditions of Lake Windermere in which species that prefer less eutrophic conditions (Atlantic salmon, brown trout, Arctic charr, minnow and bullhead) were more abundant in the mesotrophic North Basin, and species associated with eutrophic conditions (roach, tench, rudd, bream and European eel) were more common in the eutrophic South Basin (Hänfling et al. 2016). Furthermore, both of the diadromous fish lakes (Community 2) contained a different fauna including various combinations of mostly diadromous species such as European eel, three-spined stickleback, nine-spined stickleback and brown trout. Alongside these, several coarse fish species such as roach, perch, and rudd also occurred in these lakes. However, of greater conservation interest is the distribution of nine-spined stickleback. This species occurs only locally in Wales, mainly at lowland sites close to the sea (Davies et al. 2004; Hatton-Ellis 2005).

## Conclusions

In this study, we have extended the geographical, ecological and taxonomic extent of the lake fish eDNA-based metabarcoding dataset, including 14 UK lakes with well described fish faunas. Four eDNA communities are characterised, which is consistent with earlier assessments and ecological interpretations. Moreover, eDNA metabarcoding outperformed established survey techniques in terms of species detection, relative abundance using the standard five-level classification scale and characterisation of ecological fish communities, suggesting eDNA metabarcoding has great potential as fish-based assessment tool for WFD lake status assessment. Further optimisation and development such as understanding taxon-specific biases, abundance estimates and management of low confident occurrence are strongly recommended with a larger dataset, to improve the accuracy, effectiveness and applicability of WFD monitoring tool.

## Authors’ contributions

JL wrote the manuscript; BH, TWH-E, and GP conceived and designed the study; TWH-E and GP prepared historical fish data; HSK, MB, and JL carried out the fieldwork; MB prepared the sampling map; JL conducted laboratory work and performed bioinformatics analyses; JL, LLH, and BH performed the statistical analyses. All authors commented the final manuscript.

## Acknowledgments

This work was part of PhD project of JL, who was supported by the University of Hull and China Scholarship Council. This work was funded by the UK Environment Agency and Natural Resources Wales. We are particularly grateful to all the site owners and managers for granting access and the Bangor University for providing access to their laboratory facilities. Ecological Consultancy Ltd provided fishery survey data and cooperation in data collation of Cheshire meres. Robert Jaques, Hayley Watson, Mags Cousins, Peter Clabburn, Rob Evans, Sophie Gott, Emma Keenan, Ian Sims, Emma Brown, Jason Jones, Dawn Parry, Peter Shum, Harriet Johnson, Rob Donnelly and Christoph Hahn provided invaluable help during fieldwork and lab work.

## Data accessibility

Data is available from the GitHub repository (https://github.com/HullUni-bioinformatics/Li_et_al_2018_eDNA_fish_monitoring). The repository is archived with Zenodo (https://doi.org/10.5281/zenodo.1462898).

